# Coordinating Receptor Expression and Wiring Specificity in Olfactory Receptor Neurons

**DOI:** 10.1101/594895

**Authors:** Hongjie Li, Tongchao Li, Felix Horns, Jiefu Li, Qijing Xie, Chuanyun Xu, Bing Wu, Justus M. Kebschull, David Vacek, Anthony Xie, David J. Luginbuhl, Stephen R. Quake, Liqun Luo

## Abstract

The ultimate function of a neuron is determined by both its physiology and connectivity, but the transcriptional regulatory mechanisms that coordinate these two features are not well understood^1–4^. The *Drosophila* Olfactory receptor neurons (ORNs) provide an excellent system to investigate this question. As in mammals^5^, each *Drosophila* ORN class is defined by the expression of a single olfactory receptor or a unique combination thereof, which determines their odor responses, and by the single glomerulus to which their axons target, which determines how sensory signals are represented in the brain^6–10^. In mammals, the coordination of olfactory receptor expression and wiring specificity is accomplished in part by olfactory receptors themselves regulating ORN wiring specificity^11–13^. However, *Drosophila* olfactory receptors do not instruct axon targeting^6, 14^, raising the question as to how receptor expression and wiring specificity are coordinated. Using single-cell RNA-sequencing and genetic analysis, we identified 33 transcriptomic clusters for fly ORNs. We unambiguously mapped 17 to glomerular classes, demonstrating that transcriptomic clusters correspond well with anatomically and physiologically defined ORN classes. We found that each ORN expresses ~150 transcription factors (TFs), and identified a master TF that regulates both olfactory receptor expression and wiring specificity. A second TF plays distinct roles, regulating only receptor expression in one class and only wiring in another. Thus, fly ORNs utilize diverse transcriptional strategies to coordinate physiology and connectivity.

---

Single-cell RNA-sequencing (scRNA-seq) has been used to classify neurons and identify markers for specific neuronal types in diverse organisms and brain regions^15–21^. Using a recently developed scRNA-seq protocol for *Drosophila* brain cells^19^, we profiled *Drosophila* ORNs at 42–48 hours after puparium formation (hereafter “48h APF”) when ORNs are completing their axon targeting^22^ and a subset of olfactory receptors begins to be expressed^6, 8, 14^ (Fig. 1a). Briefly, the third antennal segments (which contain cell bodies of 44 out of 50 ORN classes) of transgenic flies expressing mCD8:GFP in some or all ORN classes were manually dissected, single-cell suspensions were prepared and sorted based on fluorescence, and sequencing libraries of GFP+ cells were obtained using a modified SMART-seq2 protocol (**Methods**). Each cell was sequenced to a depth of ~1 million reads, resulting in ~1500 detected genes (Extended Data Fig. 1a). After filtering cells for expression of mCD8:GFP and five neuronal markers (Extended Data Fig. 1b), 1016 high-quality ORNs were further analyzed. We did not detect expression of any markers specific to auditory neurons housed in the adjacent second antennal segment (Extended Data Fig. 1c), validating our dissection accuracy.

**Figure 1.**
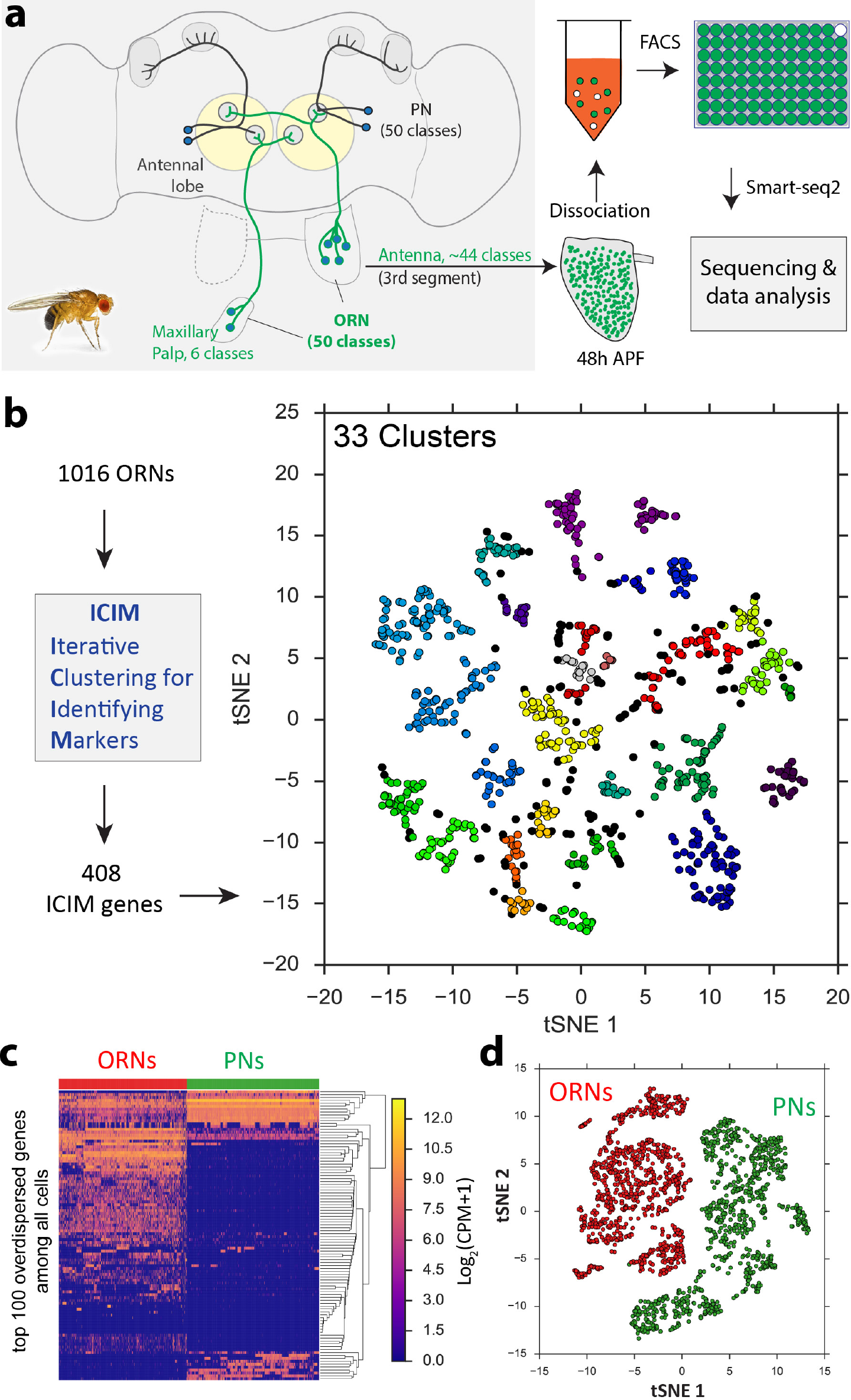
Single-cell RNA-seq for *Drosophila* olfactory receptor neurons (ORNs) at 48h APF. **a**, Schematic of fly olfactory system and single-cell RNA-seq workflow. 50 classes of ORNs (44 from the third segment of the antenna and 6 from the maxillary palp) send axons to the antennal lobe and form stereotypical one-to-one connections with 50 classes of projection neurons (PNs). The third segments of pupal antennae were manually dissected, GFP+ ORNs were sorted into 96-well plates using fluorescence-activated cell sorting (FACS), and the Smart-seq2 protocol was used for library preparation and sequencing^48^. **b**, Visualization of ORN transcriptomic clusters using t-distributed Stochastic Neighbor Embedding (tSNE) plot based on 408 genes identified by ICIM. Each dot is a cell. 1016 ORNs at 48h APF (908 from pan-ORN *nSyb-GAL4*, 63 from *85A10-GAL4*, and 45 from *AM29-GAL4*) form 33 distinct clusters. Black dots are cells that could not be assigned to any cluster. **c**, Hierarchical heat map showing clear separation of 1016 ORNs (red) and 946 PNs (green) using top 100 overdispersed genes that were identified across all cells. Each column is one cell and each row is one gene. ORNs are from 48h APF. PNs are from 24h APF^19^. Expression levels are indicated by the color bar (CPM, counts per million sequence reads). Cells (columns) and genes (rows) are ordered using hierarchical clustering. **d**, Visualization of ORNs (red) and PNs (green) using principal component analysis followed by tSNE plot using top 500 overdispersed genes. Each dot is a cell.

We sought to identify ORN types by clustering these 1016 ORNs based on transcriptomic identity. Using ICIM (Iterative Clustering for Identifying Markers), an unsupervised machine-learning algorithm^19^, we identified 408 highly informative genes for dimensionality reduction using Principal Component Analysis (PCA) and t-distributed Stochastic Neighbor Embedding (tSNE) (**Methods**). Hierarchical density-based unbiased clustering in the tSNE space revealed 33 distinct clusters from 1016 ORNs (Fig. 1b). No batch effects were evident, and sequencing depth did not drive clustering (Extended Data Fig. 1d, e).

*Drosophila* ORNs and projection neurons (PNs) are synaptic partners in the antennal lobe. We compared their transcriptomic differences using the 1016 ORNs here with 946 PNs sequenced previously^19^. Using either highly variable (overdispersed) genes across all cells, or differentially expressed genes between ORNs and PNs, ORNs and PNs were readily separated into two distinct groups (Fig. 1c, d; Extended Data Fig. 2a). We further identified several ORN- and PN-specific genes, including widely used ORN and PN markers (Extended Data Fig. 2b). Expression of *NompB*, a newly-identified ORN-specific genes, was validated using *T2A-GAL4* inserted into the *NompB* endogenous locus (Extended Data Fig. 2c)^23^.

We next employed three strategies to map transcriptomic clusters to anatomically- and functionally-defined glomerular classes of ORNs. First, we used *AM29-GAL4*^24^ to label two ORN classes (DL4 and DM6) and *85A10-GAL4*^25^ to label five ORN classes (DA1, DL3, VA1d, VA1v, and DC3), and sequenced these cells at 48h APF (Fig. 2a). To restrict GAL4 expression to ORNs, we utilized an intersectional strategy by combining GAL4 drivers with *ey-Flp* and *UAS-FRT-STOP-FRT-mCD8:GFP*^26^. As expected, *AM29+* and *85A10+* ORNs mapped to two and five different ORN clusters, respectively (Fig. 2b). Further characterization revealed the one-to-one correspondence between transcriptomic clusters and glomerular classes (see below). Second, we used *fruitless* (*fru*) expression, which is limited to three ORN classes, DA1, VA1v, and VL2a^27^, to confirm and decode three more clusters (Fig. 2c). Third, we used olfactory receptor expression to map transcriptomic clusters to glomerular classes, based on previously established correspondences between olfactory receptor expression and glomerular targets^28–30^. We systematically assessed the expression of olfactory receptors, including those belonging to the families of odorant receptors (Ors), gustatory receptors (Grs), and ionotropic receptors (Irs)^6–8, 31, 32^. Excluding co-receptors expressed in multiple adult ORN classes (e.g., *Orco* and *Ir25a*)^33, 34^, we found that 37% ORNs expressed olfactory receptors at 48h APF (Fig. 2d), consistent with the previous finding that olfactory receptors are gradually turned on in the 2^nd^ half of the pupal stage^8^. We found that receptor expression itself did not drive the clustering (Extended Data Fig. 3). Thus, the expression of olfactory receptors allowed us to decode the glomerular identity of 14 transcriptomic clusters (Fig. 2e; Extended Data Fig. 4, 5a–f).

**Figure 2.**
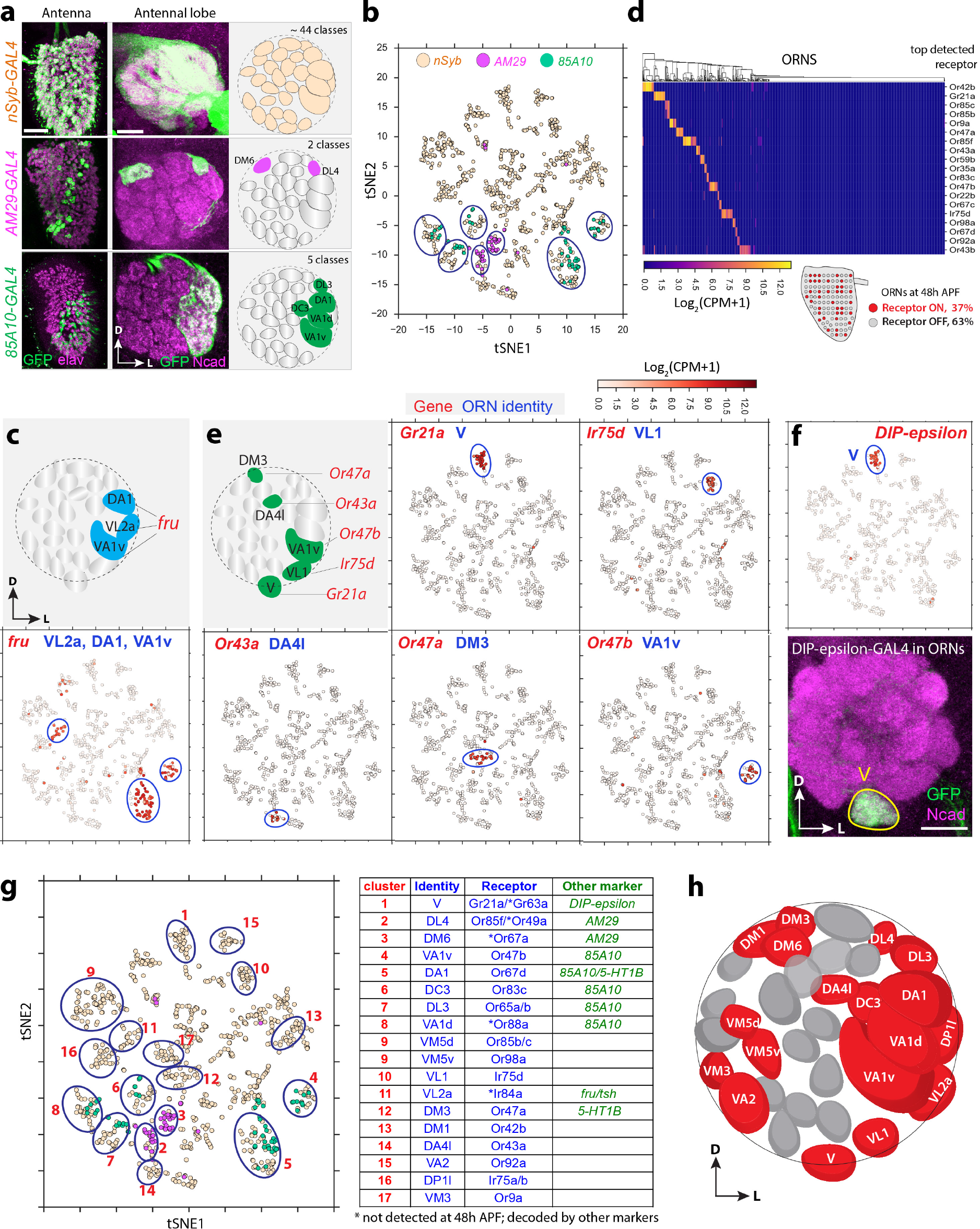
Mapping transcriptomic clusters to ORN classes. **a**, Three different drivers were used to label ORNs for single-cell RNA-seq: *nSyb-GAL4* for all ORNs, *AM29-GAL4* and *85A10-GAL4* for two and five specific ORN classes, respectively. Confocal images showing expression patterns of three drivers in both antenna and antennal lobe from 48h APF. All drivers were crossed with *ey-Flp;UAS-FRT-STOP-FRT-mCD8:GFP* to limit the GFP expression to cells in the antenna. Elav staining labels neuronal nuclei, and N-cadherin (Ncad) staining labels neuropil. Scale, 20 μm. D, dorsal; L, lateral. **b,** Visualization of *nSyb+, AM29+*, and *85A10+* ORNs using tSNE plot as in Fig. 1b. Cells are colored according to different drivers. *AM29+* cells (magenta) map to two clusters, and *85A10+* cells (green) map to five clusters (circled). Note that several individual cells from both drivers fall into other clusters, likely due to stochastic sparse labeling of these two drivers in other ORNs beyond the 7 ORN classes. **c,** Top, schematic of the fly antennal lobe showing that *fruitless* (*fru*) is expressed in three ORN classes^27^. Bottom, tSNE plots showing that *fru* expression is mostly restricted to three clusters (outlined). **d,** Heat maps showing top 19 detected olfactory receptor genes in 1016 ORNs. Each column is an individual ORN from 48h APF. 37% of ORNs show receptor expression at this stage. Cells are ordered using hierarchical clustering. **e,** Schematic of the fly antennal lobe showing selected examples of receptor expression in distinct ORN classes. tSNE plots showing expression of selected receptors. Genes and corresponding ORN classes are indicated. **f,** Validation of the decoded cluster that was mapped to V. Intersecting *ey-Flp* with *DIP-epsilon-T2A-GAL4* specifically labels V ORNs (GFP, green) at 48h APF. N-cadherin (Ncad) staining labels neuropil. Scale, 20 μm. D, dorsal; L, lateral. **g,** Summary of 17 transcriptomic clusters of ORNs that have been mapped to glomerular classes. The table on the right shows cluster identities and their corresponding receptors. Note that cluster 9 was mapped to two ORN classes VM5d and VM5v, and current analysis cannot unambiguously distinguish them. **h,** Schematic of the fly antennal lobe showing decoded ORN classes. D, dorsal; L, lateral. Expression levels are indicated by the color bar (CPM, counts per million). All tSNE plots use the same scale as Fig. 2b.

These strategies gave congruent results when mapping the same transcriptomic clusters. We further validated our cluster identity assignments using *T2A-GAL4* drivers^23^ from several genes including *DIP-epsilon*, *tsh*, and *5-HT1B*. In all cases, the GAL4 expression patterns matched their cluster identities (Fig. 2f; Extended Data Fig. 5g, h). In summary, we mapped 17 transcriptomic clusters to 18 different ORN glomerular classes (Fig. 2g, h; cluster #9 was mapped to two ORN classes, VM5d and VM5v), indicating that transcriptomic clusters correspond well with anatomically and physiologically defined ORN classes. Our annotated dataset provides a valuable resource for studying development and function of individual ORN classes.

We next investigated the mechanisms by which transcription factors (TFs) regulate olfactory receptor expression and wiring specificity in transcriptomically-identified ORN classes. In principle, three types of TFs may exist: TF-wiring (TFw) regulates only wiring specificity; TF-receptor (TFr) regulates only receptor expression; TF-master (TFm) regulates both (Fig. 3a). We first characterized TF expression at the levels of cells and clusters. We defined a TF as expressed in a cell if it reaches a threshold of 7 counts per million reads (CPM), or Log_2_ (CPM+1) ≥ 3^19^. Of the 1045 TFs in the fly genome^35^, 899 were expressed in at least one ORN, and on average about 150 TFs were detected in individual ORNs (Fig. 3b; Extended Data Fig. 6a). At the level of transcriptomic clusters, 423 TFs were detected in one or more clusters (where expression in a cluster was defined as ≥ 30% of cells in the cluster expressing the TF; Fig. 3c). These 423 TFs exhibited a wide range of expression sparsity, from a single cluster to all clusters (Fig. 3c, d). We tested several levels of thresholds that define a positive cell and percentages of cells that define a positive cluster, and obtained similar results (Extended Data Fig. 6b, c).

**Figure 3.**
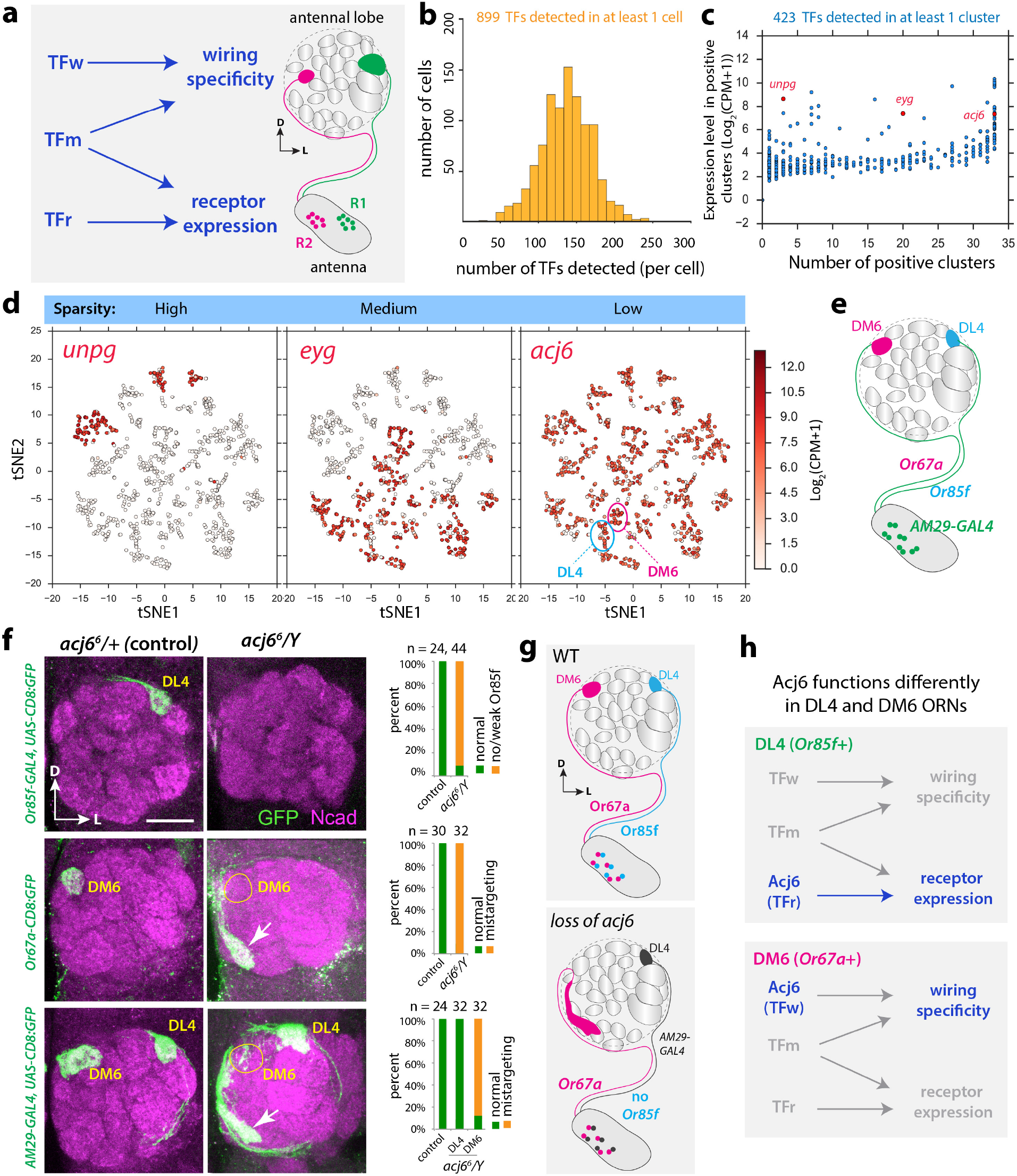
Transcription factor expression in ORNs and role of *acj6* in olfactory receptor expression and axon targeting. **a,** Schematic showing strategies for transcriptional regulation of olfactory receptor expression and wiring specificity in fly ORNs. Each ORN class expresses a unique olfactory receptor (or unique combination of receptors in several cases), and sends their axons to a specific glomerulus in the antennal lobe. Three kinds of transcription factors (TFs) may regulate these two events: TF-wiring (TFw) regulates wiring specificity, TF-receptor (TFr) regulates receptor expression, and TF-master (TFm) regulates both. **b,** Distributions of the number of TFs detected per ORN at the level of Log_2_(CPM+1) ≥ 3, or CPM ≥ 7. **c,** Sparsity and expression level of the TFs among ORN transcriptomic clusters. Each dot is one TF. A positive cluster is defined as more than 30% cells in the cluster expressing the TF at the level of Log_2_(CPM+1) ≥ 3. Highlighted in red are three example genes, *unpg*, *eyg*, and *acj6*, with high mean expression levels but different sparsity. **d,** tSNE plots showing expression of *unpg, eyg,* and *acj6*. Expression levels are indicated by the color bar (CPM, counts per million). In the *acj6* tSNE plot, two clusters corresponding to DL4 and DM6 are indicated. **e,** Schematic showing that *AM29-GAL*4 labels two ORN classes DL4 and DM6, which express *Or85f* and *Or67a*, respectively. **f,** Confocal images of adult antennal lobes showing ORN axon targeting in heterozygous control (left) and hemizygous mutant (right). In control, *Or85f*+ ORNs target to DL4, *Or67a*+ ORNs target to DM6, and *AM29*+ ORNs target to both DL4 and DM6. In mutant, *Or85f* expression is lost, but DL4 ORN axons still target normally as shown by *AM29-GAL4*; *Or67a* expression is normal, but *Or67a*+ ORNs show mistargeting (arrow in the middle panel), as confirmed by *AM29-GAL4* (arrows in the bottom panel). All images are confocal z-stacks covering the region of targeted glomeruli. N-cadherin (Ncad) staining labels neuropil. Scale, 20 μm. D, dorsal; L, lateral. Quantifications are shown on the right. Antennal lobe numbers (n) are indicated. **g,** Schematic summary of data in (**f**). **h,** In DL4 ORNs, *ajc6* is only required for receptor expression, but not for wiring specificity. In DM6 ORNs, *ajc6* is only required for wiring specificity, but not for receptor expression.

Previous studies have identified several TFs that regulate *Drosophila* olfactory receptor expression or wiring specificity^36–39^. However, in all cases, these two processes have been investigated separately because of a lack of appropriate genetic tools. When a TF is required for olfactory receptor expression in an ORN class, it has not been possible to investigate whether this TF also regulates wiring because the ORN-specific marker for examining glomerular targeting is usually a transgene driven by the promoter of the olfactory receptor itself. We overcame this obstacle by using two ORN classes—DL4 (*Or85f+)* and DM6 (*Or67a+*)—that can be independently labeled by *AM29-GAL4* (Fig. 3e).

We first focused on a POU-domain transcription factor, *abnormal chemosensory jump 6* (*acj6*), which is expressed in most antennal ORNs and regulates receptor expression in some ORN classes and axon targeting in other ORN classes^36, 37, 39^. However, it is unclear if *acj6* regulates both wiring specificity and receptor expression in the same ORN class. Our sequencing data showed that *acj6* was expressed in all ORN clusters, including DL4 and DM6 (Fig. 3c, d). Using an *acj6* null allele (*acj6^6^*)^36^, we first confirmed that *acj6* is required for the expression of *Or85f* in DL4 ORNs, but not required for the expression of *Or67a* in DM6 ORNs (Fig. 3f), as previously reported^39^. Using *AM29-GAL4* to label the axons of DL4 and DM6 ORNs, we found that DM6 ORNs showed highly penetrant mistargeting phenotypes, while DL4 ORNs still targeted to the correct glomerulus (Fig. 3f, g). Thus, *acj6* is not a TFm in DL4 or DM6 ORNs. Rather, it is a TFr in DL4 ORNs, but a TFw in DM6 ORNs (Fig. 3h).

The expression specificity of olfactory receptors or axon guidance molecules could be controlled by at least two distinct mechanisms in terms of TF expression: 1) the expression of the regulatory TF is restricted to certain ORN classes; 2) the TF is widely expressed, but its activity is restricted to specific ORN classes by specific expression of other transcriptional co-activators or repressors. To date, TFs known to control olfactory receptor expression are mostly widely expressed, supporting the combinatorial activation/repression model^6, 38, 39^. To test whether class-restricted TF expression also contributes to olfactory receptor expression and wiring specificity, we identified several TFs expressed at high levels in only a few transcriptomic clusters (Fig. 3c, d). We performed an RNAi screen on 25 candidate TFs, using a pan-ORN driver *peb-GAL4*^40^ to knockdown candidate genes, and monitored the expression of olfactory receptors using receptor promoter-driven reporters. This screen revealed the role of *unplugged* (*unpg*), which encodes a homeobox transcription factor, in regulating olfactory receptor expression.

*unpg* was previously identified as regulating tracheal branch formation^41^ and being a marker for a specific neuroblast sub-lineage^42^. We found *unpg* expression in three transcriptomic clusters mapped to four ORN classes: V (*Gr21a+*), VA2 (*Or92a+*), VM5d (*Or85b+*), and VM5v (*Or98a*+) (Fig. 4a, b; transcriptomes of VM5d and VM5v could not be unambiguously distinguished, as shown in Fig. 2g). Because the *Or85b* reporter was barely detectable in control flies (data not shown), we focused our subsequent analysis on the other three ORN classes. We found that expression of all three receptors, *Or92a*, *Or98a*, and *Gr21a*, was lost when *unpg* was knocked down in ORNs using RNAi (Fig. 4c, d; Extended Data Fig. 7a). By contrast, receptor expression in four *unpg*-negative ORN classes (*Or42b, Or10a, Or88a,* and *Or47b*) was unaffected (Fig. 4d; Extended Data Fig. 7b, c), suggesting that *unpg* is specifically required for olfactory receptor expression in *unpg*-positive ORNs. Since *Or42b+* and *Or10a+* ORNs (both *unpg*-negative) are from the same ab1 sensilla in the antenna as *Or92a+* and *Gr21a+* ORNs (*unpg*-positive) (see Fig. 4g), these data suggest that the loss of *Or92a* and *Gr21a* in *unpg-RNAi* flies was not due to gross developmental defects of the ab1 sensilla.

**Figure 4.**
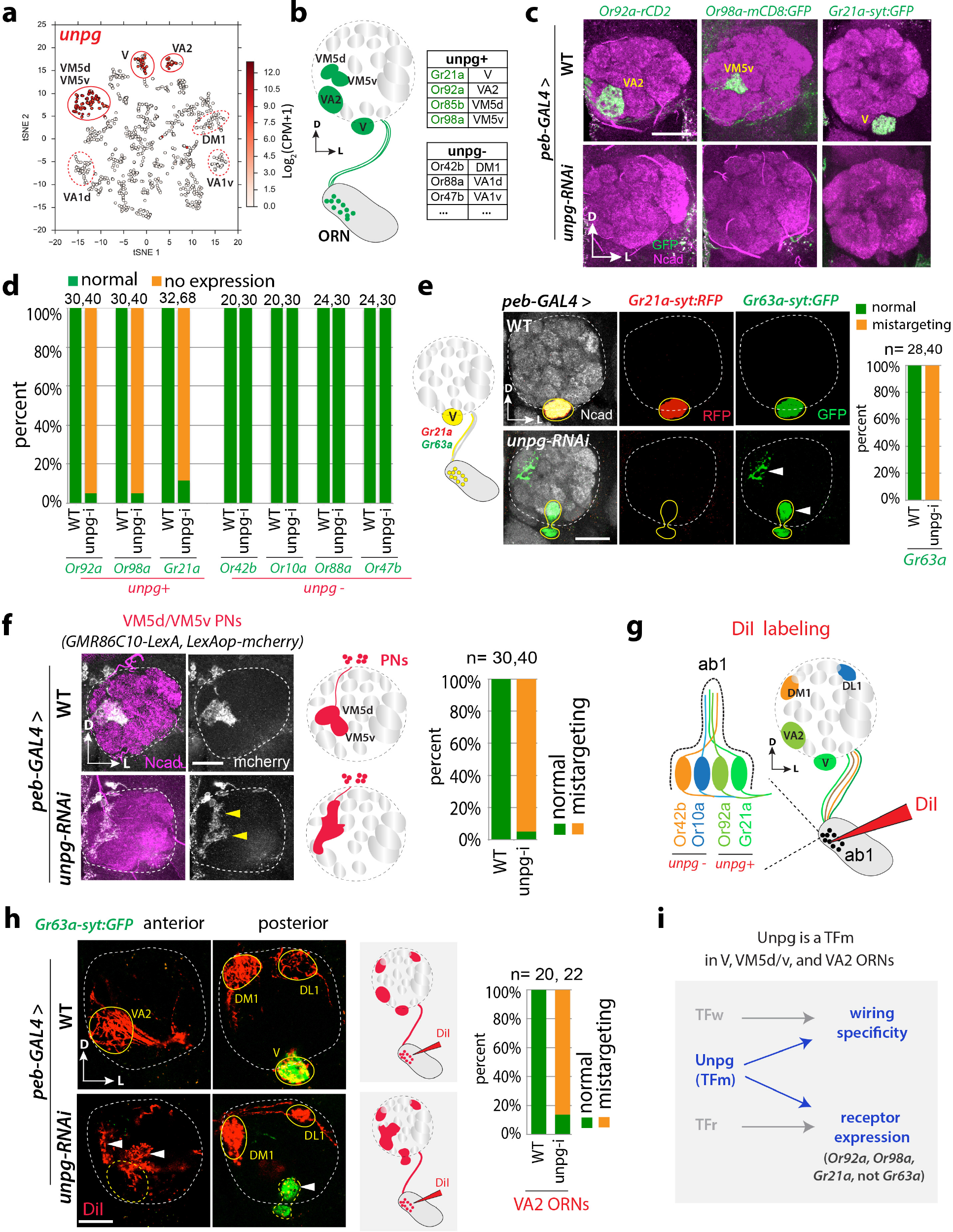
*unpg* controls receptor expression and wiring specificity in *unpg+* ORN classes. **a,** tSNE plot showing that *unpg* is expressed in three clusters mapped to V, VA2, and VM5d/VM5v (solid outline). Three *unpg*-negative clusters are indicated (dashed outline). **b,** Schematic summarizing receptor expression and glomerular targets of all *unpg*-positive and some *unpg*-negative ORN classes. **c,** Confocal images from adult flies showing olfactory receptor expression in control (WT) and *unpg-RNAi* flies. *peb-GAL4;UAS-dcr2* was crossed with either *w*^*1118*^ (WT) or *unpg-RNAi*, and olfactory receptor promoter-driven reporters are used to monitor receptor expression. In WT flies, *Or92a, Or98a,* and *Gr21a* are expressed normally. In *unpg-RNAi* flies, the expression of all three receptors is lost. **d,** Quantification for the expression of different receptors, including three *unpg*-positive ones in (**c**) and four *unpg*-negative ones. All four *unpg*-negative receptors show normal expression in *unpg-RNAi* flies. Antennal lobe numbers (n) are indicated on top. **e,** *Gr63a* and *Gr21a* are co-expressed in V ORNs. For confocal images, *peb-GAL4;UAS-dcr2* was crossed with either *w*^*1118*^ (WT) or *unpg-RNAi*, and *Gr21a-syt:RFP* and *Gr63a-syt:GFP* were simultaneously used to label V ORNs. In *unpg-RNAi* flies, *Gr21a* expression is lost consistently, and *Gr63a*+ axons show stereotyped mistargeting (arrowheads). See more images in Extended Data Fig. 7d. Quantifications are shown on the right; antennal lobe numbers (n) are indicated. **f,** VM5d and VM5v PNs are labeled by *GMR86C10-lexA;lexAop-mCherry*, and their dendrite targeting are monitored in WT and *unpg-RNAi* (crossed with *peb-GAL4;UAS-dcr2)* flies. In WT flies, labeled PNs send dendrites to VM5d and VM5v glomeruli with clear boundary. In *unpg-RNAi* flies, these PNs send dendrites to diffuse regions with fuzzy boundary (arrowheads). Schematic VM5d and VM5v PNs are shown on the right top. See more images in Extended Data Fig. 7e. Quantifications are shown on the right; antennal lobe numbers(n) are indicated. **g,** DiI labeling for four ORN classes from the same ab1 sensilla in the antenna: *Or42b* and *Or10a* (*unpg*-negative), as well as *Gr21a* and *Or92a* (*unpg*-positive). **h,** For DiI labeling, *Gr63a-syt:GFP* is used to monitor V ORNs. In WT flies, DiI labels VA2 in an anterior section of the antennal lobe, and DM1, DL1 and V in a posterior section. In *unpg-RNAi* flies, DiI labels a diffuse area (arrowheads) around VA2 in the anterior section, labels DL1 and DM1 normally, and co-labels *Gr63a*-posistive V ORNs, which show mistargeting phenotypes. Schematic on the right summarizes DiI labeling in WT and *unpg-RNAi* flies. See more images in Extended Data Fig. 7f. Quantifications are shown on the right; antennal lobe numbers (n) are indicated. **i,** *Unpg* is a master transcription factor (TFm) in V, VM5d/VM5v, and VA2 ORNs, regulating both receptor expression and wiring specificity. Note that in V ORNs, *unpg* is required for the expression of *Gr21a*, but not for *Gr63a*, suggesting that these two co-receptors are regulated by different mechanisms. All confocal images are from adult flies, and are z-stacks covering the targeted glomeruli. N-cadherin (Ncad) staining labels neuropil (**d, f**, and **g**). Scale, 20 μm. D, dorsal; L, lateral.

We next investigated if *unpg* also regulate axon targeting in *unpg*-positive ORNs using markers independent of their receptors. V ORNs co-express a pair of gustatory receptors, *Gr21a* and *Gr63a*, for carbon dioxide sensing^43, 44^. We utilized *Gr63a-syt:GFP* as an independent marker for V ORNs and found it was normally expressed in *unpg-RNAi* flies. Interestingly, *Gr63a+* axons of V ORNs showed stereotyped mistargeting in all *unpg-RNAi* flies (Fig. 4e; Extended Data Fig. 7d). Thus, *unpg* regulates both *Gr21a* expression and axon targeting in V ORNs.

Since there are no independent markers for VM5d, VM5v, and VA2 ORNs, we utilized two alternative strategies to study their axon targeting. We identified a Janelia-GAL4 driver^25^, *GMR86C10-GAL4*, that specifically labels VM5d and VM5v PNs, and generated a *GMR86C10-LexA* driver to label these PNs independent of the GAL4/UAS system. Since ORN axon mistargeting during development can lead to dendrite mistargeting of partner PNs^45, 46^, we reasoned that we might observe mistargeting of VM5d and VM5v PNs if knocking down *unpg* in VM5d and VM5v ORNs causes targeting defects. This was indeed the case (Fig. 4f; Extended Data Fig. 7e), suggesting that *unpg* also regulates VM5d and VM5v ORN axon targeting.

Finally, we utilized DiI-mediated anterograde tracing to examine axon targeting of VA2 (*Or92a+*) ORNs. Each ab1 sensillum houses four ORNs: *Or42b+* and *Or10a*+ ORNs are *unpg*-negative, while *Gr21a*+ and *Or92a*+ ORNs are *unpg*-positive (Fig. 4g)^28^. As expected, knockdown of *unpg* did not affect targeting of *Or42b*+ or *Or10a*+ ORNs to DM1 or DL1 glomeruli (Extended Data Fig. 7c). By applying DiI to a small subset of ab1 sensilla to initiate anterograde tracing, we found a stereotyped mistargeting zone dorsal to the V glomerulus coinciding with mistargeted V ORN axons labeled by *Gr63a-syt:GFP.* Notably, diffuse mistargeting was also observed dorsal to the VA2 glomerulus in most *unpg-RNAi* flies (Fig. 4g, h; Extended Data Fig. 7f), suggesting that *unpg* also regulates axon targeting of VA2 ORNs. Taken together, our data suggest that *unpg* is a TF-master (TFm) in all examined *unpg*-positive ORN classes, where it controls both olfactory receptor expression and wiring specificity (Fig. 4i).

Our understanding of how developing neurons coordinately regulate physiological properties and connectivity is limited to few examples. In the mouse olfactory system, an ORN’s olfactory receptor, which specifies its physiological responsivity, is also utilized to instruct axon targeting^11–13^. In *Drosophila* R8 photoreceptors, the TF *Senseless* promotes the expression of both an R8-specific rhodopsin and a transmembrane protein, Capricious, that regulates axon targeting^47^. Here, we found that even in the same group of neurons (*Drosophila* ORNs), the coordination of these two features uses diverse transcriptional strategies in different ORN classes (Extended Data Fig. 7g). On one hand, a broadly expressed TF, Acj6, regulates receptor expression but not wiring in one ORN class, and wiring but not receptor expression in a second class. These data suggest that Acj6 acts in a combinatorial manner with other transcriptional activators and repressors in an ORN class- and target (olfactory receptors vs wiring molecules)-specific manner. On the other hand, a more specifically expressed TF, Unpg, regulates both receptor expression and wiring specificity in all classes that express Unpg. Future studies to identify transcriptional targets of these TFs, and to investigate how different TFs interact with each other, will provide further insight into how these diverse transcriptional regulatory strategies are executed. Given that each ORN expresses ~150 TFs, it is remarkable that disruption of a single TF can result in profound disruption of receptor expression and wiring specificity, two of the most fundamental properties of a sensory neuron.

In conclusion, scRNA-seq in developing *Drosophila* ORNs enabled us to map transcriptomic identity to anatomical and physiological identity for 17 ORN classes. This reinforced the idea that neuronal transcriptome identity corresponds well with anatomical and physiological identities defined by connectivity and function in well-defined neuronal types^17, 19^. Our transcriptome map for glomerular classes provides a valuable resource for future investigation into the development and physiology of the *Drosophila* olfactory system. Finally, our scRNA-seq analysis has revealed, to our knowledge, the first transcription factor that coordinates olfactory receptor expression and neuronal wiring in the fly olfactory system, highlighting the power of scRNA-seq for investigating mechanisms that control neural development.

## Acknowledgements

We thank H. Bellen, L. Zipursky, I. Grunwald Kadow, J. Simpson, Bloomington and Vienna Stock Centers for reagents; J. Lui and E. Richman for discussions; N. Neff, J. Okamoto, B. Jones and Chan-Zuckerberg BioHub for assistance with sequencing; Y. Ge for assistance on fly work; and A. Shuster, J. Lui, and D. Pederick for comments on the manuscript. H.L. is a Stanford Neuroscience Institute Interdisciplinary Postdoctoral Scholar, F.H. acknowledges support from the National Science Foundation Graduate Research Fellowship, J.L. thanks Genentech Foundation Predoctoral and Vanessa Kong Kerzner Graduate Fellowships. S.R.Q. is a Chan Zuckerberg Investigator, and L.L. is an HHMI Investigator. This work was supported by NIH grant R01-DC005982 (to L.L.).

## Author contributions

H.L., T.L., and L.L. designed experiments with support from F.H. and S.R.Q. H.L. and T.L collected samples and performed scRNA-seq. H.L. analyzed data with help from F.H., and J.M.K. T.L. performed DiI labeling experiment and H.L. performed all other fly experiments with help from T.L., J.L., Q.X., C.X., B.W., D.V., A.X., and D.J.L. Q.X. performed qRT-PCR. H.L. and L.L. wrote the paper with input from T.L., F.H., and S.R.Q.

## Methods

### Fly stocks

The following fly lines were used in this study. *nSyb-GAL4* (Bloomington *Drosophila* Stock Center, BDSC #51635); *act5C-GAL4* (BDSC #3954); *AM29-GAL4*^24^; *GMR86C10-GAL4* and *GMR85A10-GAL4*^25^; *NompB-T2A-GAL4* (BDSC #76632); *DIP-epsilon-T2A-GAL4* (BDSC #67502); *5-HT1B-T2A-GAL4* (BDSC #76668); *tsh-T2A-GAL4*^23^; *acj6*^6, 36^; *Mz19-QF*^45^; *unpg-RNAi* (VDRC stock #107638); *peb-GAL4*^40^; *GH146-Flp*^49^; *ey-Flp*^50^; *UAS-STOP-mCD8:GFP*^26^; *Or-GAL4* and *Or-mCD8:GFP*^28, 37^; *Or85f-GAL4, Or42b-mCD8:GFP, Or67a-mCD8:GFP, Or98a-mCD8:GFP; Or92a-rCD2*^51^; *Or47b-rCD2*^52^; *Or88a-mtdT*^46^; *Or10a-LexA*^29^; *Gr21a-syt:GFP* and *Gr63a-syt:GFP*^43^.

### Immunostaining

Tissue dissection and immunostaining were performed following previously described methods^53^. Briefly, fly pupal and adult brains were dissected in 1x PBS and then fixed in 4% paraformaldehyde (20% paraformaldehyde diluted in PBS with 0.015% Triton X-100) for 20 min at room temperature. Fixed brains were washed three times with PBST (PBS with 0.3% Triton X-100) and incubated in PBST twice for 20 min. The samples were incubated in blocking buffer (5% normal goat serum in PBST) for 30 min at room temperature or overnight at 4°C. Then, primary antibodies diluted in blocking buffer were applied and samples were incubated for 24–48 h at 4°C. Then, samples were washed using PBST for 20 min twice, and secondary antibodies diluted in blocking buffer were applied and samples were incubated in dark for more than 24 h at 4°C. Samples were washed in PBST for 20 min twice and mounting solution (Slow Fade Gold) was added. Samples were left in mounting solution for at least 1 h before mounting them onto glass slides. All wash steps were performed at room temperature. Primary antibodies used in this study include rat anti-DNcad (DN-Ex #8; 1:40; DSHB), chicken anti-GFP (1:1000; Aves Labs), rabbit anti-DsRed (1:250; Clontech), mouse anti-ratCD2 (OX-34; 1:200; AbD Serotec). Secondary antibodies were raised in goat or donkey against rabbit, mouse, rat, and chicken antisera (Jackson Immunoresearch), conjugated to Alexa 405, 488, FITC, Cy3, Cy5, or Alexa 647.

### Confocal imaging

All confocal images were taken through a Z-stack scan from most anterior to the most posterior of the antenna or antennal lobe using the Zeiss LSM 780 system. Then images were processed with ImageJ and Adobe Illustrator. For quantification in Figs. 3 and 4, at least 10 flies (20 antennal lobes) were used.

### DiI labeling

DiI (Sigma 468495) in saturated DMSO solution was applied to the medioproximal corner of the third antennal segment of adult flies, where the ab1 sensilla are located, through glass micropipette using a micromanipulator. Flies were recovered for 24h after labeling before their brains were dissected, mounted in 30% sucrose, and imaged using a confocal microscope. Mistargeting phenotypes of both antennal lobes from each fly were quantified in WT control and *unpg-RNAi* expressing flies. Out of 11 WT flies, only 1 was labeled for more than 4 glomeruli, which is likely due to labeling of additional sensilla near ab1 during DiI application. 3 out of 14 *unpg-RNAi* expressing flies showed similar additional labeling and were excluded from quantification. In total, 10 WT and 11 *unpg-RNAi* flies (20 and 22 antennal lobes) were quantified for VA2 ORN axon targeting.

### Quantitative PCR (qPCR) for *unpg-RNAi*

Total RNA was extracted using MiniPrep kit (Zymo Research, R1054) from either *actin5C-GAL4*, *w*^*1118*^ (control) or *actin5C-GAL4;unpg-RNAi* flies at middle pupal stage (*N* = 3 replicates for each condition; 5 pupae per replicate). cDNA was synthesized using an oligo-dT primer. qPCR was performed on a Bio-Rad CFX96 detection system. *p*-value was calculated using Student’s *t* test. Relative expression was normalized to *actin5C*. Primer sequences used for qPCR were:

*actin5C* (F): 5’-CTCGCCACTTGCGTTTACAGT-3’
*actin5C* (R): 5’-TCCATATCGTCCCAGTTGGTC-3’
*unpg* pair 1 (F): 5’-CTACAACGGCGAGATGGACA-3’
*unpg* pair 1 (R): 5’-TTGGAGTTTGAGCTGGAGCC-3’
*unpg* pair 2 (F): 5’-GGAACTACAACGGCGAGATG-3’
*unpg* pair 2 (R): 5’-GATACTTCTTGGCGTGGAACT C-3’

### Single-cell RNA-seq

Single-cell RNA-seq was performed following the protocol that we developed recently ^19^. Briefly, *Drosophila* third antennal segments with mCD8:GFP-labeled cells using specific GAL4 drivers were manually dissected. Single-cell suspensions were then prepared. Single labeled cells were sorted via Fluorescence Activated Cell Sorting (FACS) into individual wells of 96-well plates containing lysis buffer using an SH800 instrument (Sony Biotechnology). Full-length poly(A)-tailed RNA was reverse-transcribed and amplified by PCR following the SMART-seq2 protocol^48^ with several modifications as below. To increase cDNA yield and detection efficiency, we increased the number of PCR cycles to 25. To reduce the amount of primer dimer PCR artifacts, we digested the reverse-transcribed first-strand cDNA using lambda exonuclease (New England Biolabs) (37°C for 30 min) prior to PCR amplification. Sequencing libraries were prepared from amplified cDNA using tagmentation (Nextera XT). Sequencing was performed using the Illumina Nextseq 500 platform with paired-end 75 bp reads.

### scRNA-seq data processing

#### Data and software availability

Sequencing data and Python code for figures are currently available upon request, and will be uploaded later to Gene Expression Omnibus and Github.

#### Sequence alignment and preprocessing

Reads were aligned to the *Drosophila melanogaster* genome (r6.10) using STAR (2.4.2a)^54^ with the ENCODE standard options, except "--outFilterScoreMinOverLread 0.4 --outFilterMatchNminOverLread 0.4 -- outFilterMismatchNmax 999 --outFilterMismatchNoverLmax 0.04". Uniquely mapped reads that overlap with genes were counted using HTSeq-count (0.7.1)^55^ with default settings except "-m intersection-strict". Cells having fewer than 300,000 uniquely mapped reads were removed. To normalize for differences in sequencing depth across individual cells, we rescaled gene counts to counts per million (CPM). All analyses were performed after converting gene counts to logarithmic space via the transformation Log_2_(CPM+1). Sequenced cells were filtered for expression of canonical neuronal genes (*elav, brp, Syt1, nSyb, CadN,* and *mCD8GFP*), retaining only those cells that expressed at least 4/6 genes at Log_2_ (CPM+1) ≥ 3.

#### PCA and tSNE

Principal component analysis (PCA) and t-distributed Stochastic Neighbor Embedding (tSNE) were used for visualizing *Drosophila* scRNA-seq data as detailed previously ^19^. Briefly, to visualize and interpret high dimensional gene expression data, we obtained two-dimensional projections of the cell population by first reducing the dimensionality of the gene expression matrix using PCA, then further reducing the dimensionality of these components using tSNE^56^. We performed PCA for Fig. 1d on a reduced gene expression matrix composed of the top 500 overdispersed genes. The top 8 principal components (8 PCs) were used. We further reduced these components using tSNE to project them into a two-dimensional space.

#### Iterative Clustering for Identifying Markers (ICIM)

We previously developed an unsupervised machine learning algorithm called ICIM to identify genes that distinguish transcriptome clusters for different fly olfactory projection neuron (PN) subtypes^19^ (available at https://github.com/felixhorns/FlyPN). We performed similar analysis for ORNs with several modifications of the adjustable parameters, including Pearson Correlation and Dropout threshold, and identified 408 ICIM genes, which we used for further dimensionality reduction by PCA. The top 18 PCs were further reduced to two-dimensions using tSNE. HDBSCAN, a hierarchical density-based unbiased clustering algorithm^57^, was used to reveal clusters.

We observed that standard dimensionality reduction and clustering methods using PCA and tSNE failed to discriminate subpopulations that corresponded to known lineages and molecular features of PNs^19^ or ORNs (current study). We attributed the failure of these methods to the high degree of similarity of transcriptional states among olfactory neuron classes, which represent closely-related neurons having similar functions. Thus, olfactory neuron classes may be distinguished by a small number of genes. We developed ICIM as a strategy to identify the most informative genes for distinguishing subpopulations within a population of closely-related cells in an unbiased way.

Briefly, starting with a population of cells, we first identify the top 100 overdispersed genes. Next we expand this set of genes by finding genes whose expression profiles are strongly correlated. We also filter this set of genes by (1) removing those having <2 correlated partners to remove noisy genes, and (2) those that are expressed in >80% of cells to remove housekeeping genes. Cells are then clustered. We cut the dendrogram at the deepest branch and partition the population into two subpopulations. The same steps are then performed iteratively on each subpopulation. Iteration continues until a population cannot be split. The termination condition is defined as the minimum terminal branch length being larger than 0.2. The result of this analysis is a set of genes that discriminate subpopulations within a population, which can be used for dimensionality reduction.

#### Overdispersion analysis and differential expression analysis

Overdispersion analysis and differential expression analysis were previously described^19^. Briefly, genes that are highly variable within a population often carry important information for distinguishing cell types.

Variability of gene expression depends strongly on the mean expression level of a gene. This motivates the use of a metric called dispersion, which measures the variability of a gene’s expression level in comparison with other genes that are expressed at a similar level. Overdispersed genes are those that display higher variability than expected based on their mean expression level. To identify overdispersed genes, we binned genes into 20 bins based on their mean expression across all cells. We then calculated a log-transformed Fano factor D(x) of each gene x

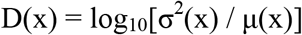

where σ^2^(x) is the variance and μ(x) is the mean of the expression level of the gene across cells.

Finally, we calculated the dispersion d(x) as the Z-score of the Fano factor within its bin

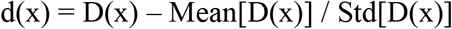

where Mean[D(x)] is the mean log-transformed Fano factor within the bin and Std[D(x)] is the standard deviation of the log-transformed Fano factor within the bin. We then rank genes by their dispersion and select the top genes for downstream analysis.

To find differentially expressed genes, we used the Mann-Whitney U test, a non-parametric test that detects differences in the level of gene expression between two populations. The Mann-Whitney U test is advantageous for this application because it makes very general assumptions: (1) observations from both groups are independent and (2) the gene expression levels are ordinal (i.e., can be ranked). Thus the test applies to distributions of gene expression levels across cells, which rarely follow a normal distribution. Using the Mann-Whitney U test, we compared the distributions of expression levels of every gene separately. P values were adjusted using the Bonferroni correction for multiple testing. Different significance thresholds for determining whether a gene is differentially expressed were used for various analyses in this work.

#### Transcription factor (TF) gene lists

To identify genes that are transcription factors (TFs), we used manually curated lists. We obtained a list of *Drosophila* TFs from the FlyTF v1 database (http://www.mrc-lmb.cam.ac.uk/genomes/FlyTF)^35^. These lists were manually curated to remove spurious annotations and redundancies according to Flybase annotation, resulting in 1045 TFs, which were used for analysis in Fig. 3 and Extended Data Fig. 6.

---

**Extended Data Figure 1.**
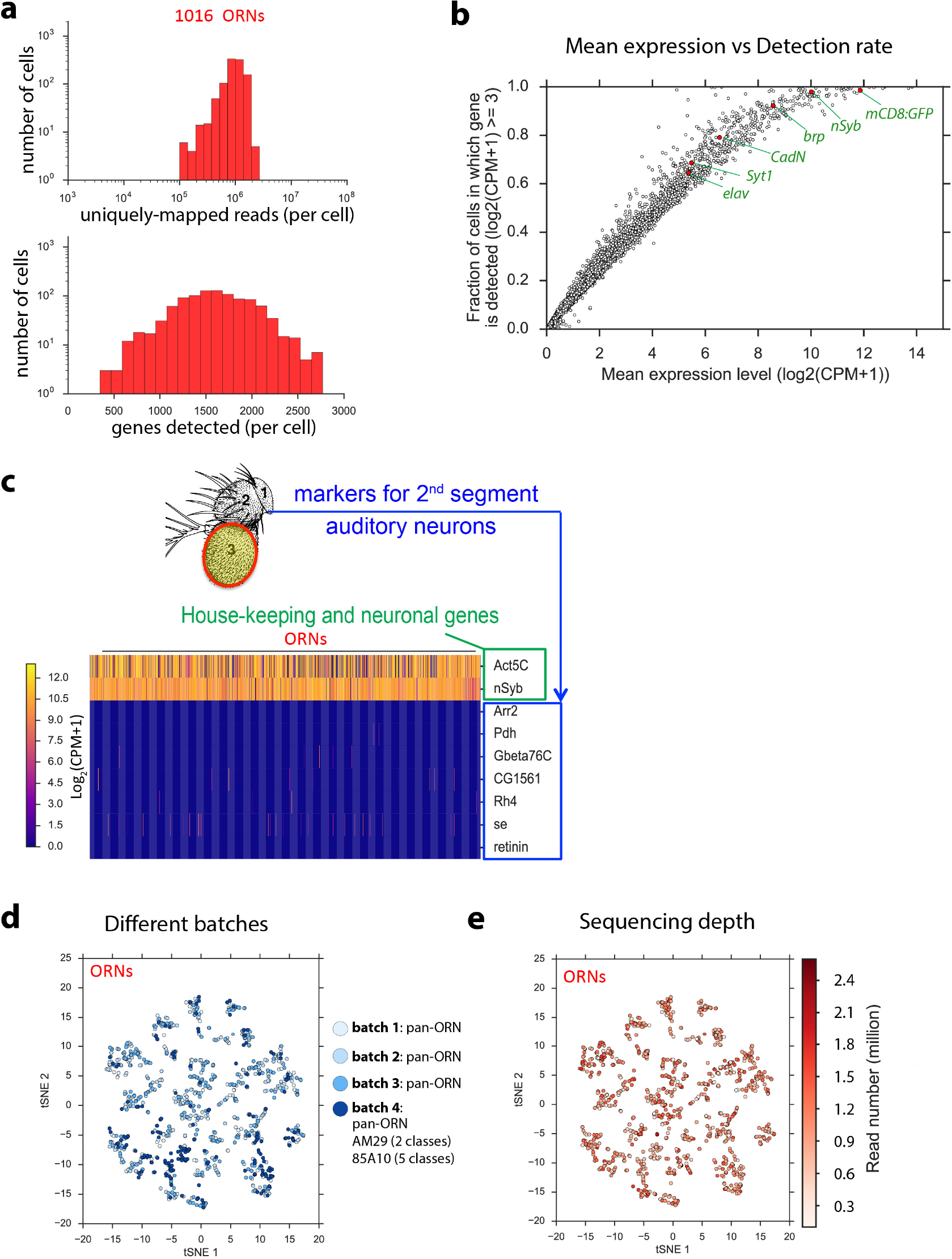
Single-cell RNA-seq for *Drosophila* olfactory receptor neurons (ORNs) at 48h APF. **a,** Distributions of uniquely mapped reads (top) and detected genes (bottom) per cell in the 1016 ORNs we analyzed. **b,** Mean expression level and detection rate of all detected genes in ORNs. Each dot is a gene. Detection is defined as Log_2_(CPM+1) ≥ 3. Detection failure events can occur because (1) the gene is not expressed in the cell, or (2) failure to detect expression of the gene despite the presence of mRNA transcripts due to technical artifact (dropouts). Thus, the fraction of detection failure events provides an upper bound on dropout rate. mCD8:GFP and the 5 neuronal markers, which were used for quality filtering, are indicated. **c,** Heat map showing expression in individual ORNs of the housekeeping gene (*actin5c)*, neuronal marker (*nSyb*), and seven genes known to be specific to auditory neurons in the 2^nd^ segment of the antenna^58^. **d,** tSNE plot showing ORNs from different batches, suggesting that there is no obvious batch effect for current clustering. Note that some ORNs from batch 4 are from two specific drivers that only label a few ORN classes, so that these ORNs are enriched in certain clusters as expected. **e,** tSNE plot showing ORNs clusters with sequencing depth indicated, suggesting that the sequencing depth does not driver current clustering.

**Extended Data Figure 2.**
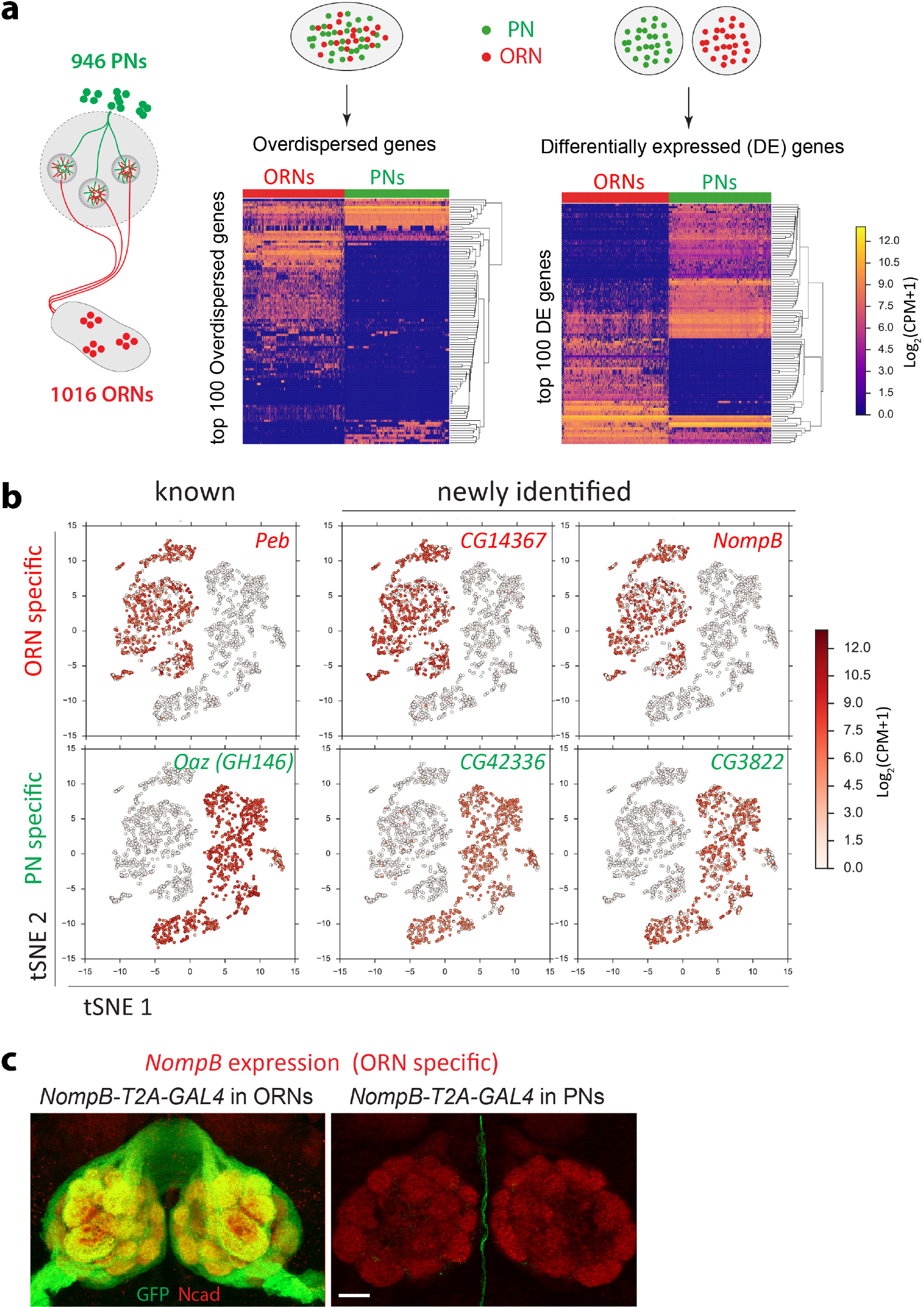
Comparison between PNs and ORNs from single-cell RNA-seq analysis. **a,** Hierarchical heat maps showing that 1016 ORNs (red) and 946 PNs (green) form distinct clusters using top 100 overdispersed genes that were identified across all cells (left heat map, same as Fig. 1c), or using top 100 differentially expressed (DE) genes that were identified by comparing ORNs with PNs (right heat map). Each column is one cell and each row is one gene. ORNs are from 48h after puparium formation (APF). PNs are from 24h APF as described in our previous study 19. Expression levels are indicated by the color bar (CPM, counts per million). Cells and genes are ordered using hierarchical clustering. **b,** tSNE plots showing expression levels of selected genes that are specific to either ORNs or PNs. *pebbled* (*peb*) and *Oaz* (where *GH146-GAL4* is inserted) are known markers for ORNs and PNs, respectively. Other four genes are examples of newly identified genes specific to ORNs or PNs. **c,** Confocal images of *Drosophila* antennal lobe at 48h APF to validate the *NompB* expression pattern as shown in (B). *NompB-T2A-GAL4* was intersected with either *ey-Flp* or *GH146-Flp* (with the *UAS-FRT-STOP-FRT-mCD8:GFP* reporter) to limit the GAL4 expression to ORNs or PNs, respectively. *NompB-T2A-GAL4* can be detected in most ORNs (top), but not in PNs (bottom), consistent with our sequencing data. N-cadherin (Ncad) staining labels neuropil. Scale, 20 μm.

**Extended Data Figure 3.**
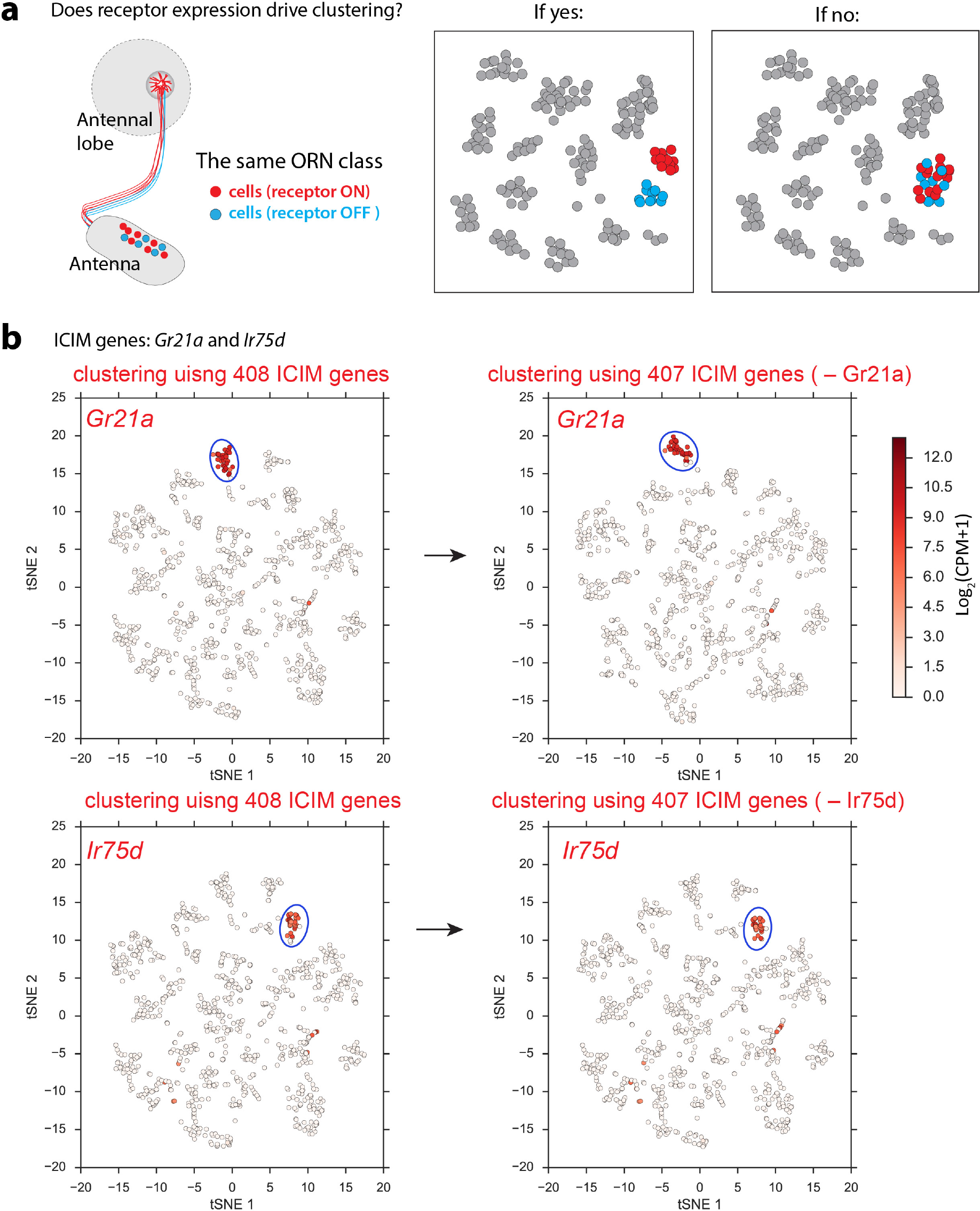
Olfactory receptor gene expression does not drive clustering. **a,** Schematics showing two alternative models. At 48h APF, olfactory receptor genes start to be expressed in some ORN classes. At this stage, within the same ORN class containing on average 30 cells in each antenna, some show receptor expression, and others do not. If the receptor expression drives clustering, the receptor-ON and receptor-OFF cells from the same ORN class would form distinct clusters (left), in which case the receptor expression is not a reliable marker to decode cluster identities. If receptor expression does no drive clustering, the receptor-ON and receptor-OFF cells from the same ORN class would form one cluster (right) due to the transcriptomic similarity without the receptor gene. In this case, receptor expression can be used to decode ORN clusters. **b,** tSNE plots using either 408 ICIM genes, or 407 genes without *Gr21a* or without *Ir75d*. There are three receptor genes in the 408 ICIM genes, *Gr21a, Ir75d*, and *Gr93a*. *Gr21a* and *Ir75d* label distinct single clusters. *Gr93a* is expressed in multiple (~10) clusters (data not shown). When *Gr21a* or *Ir75d* is removed from the ICIM genes, similar clusters are formed, and the expression of *Gr21a* or *Ir75d* is still enriched in individual clusters. Expression levels are indicated by the color bar (CPM, counts per million). These data indicate that receptor expression does not driver clustering in our analysis, as shown in the right schematic plot of Extended Data Figure 3a.

**Extended Data Figure 4.**
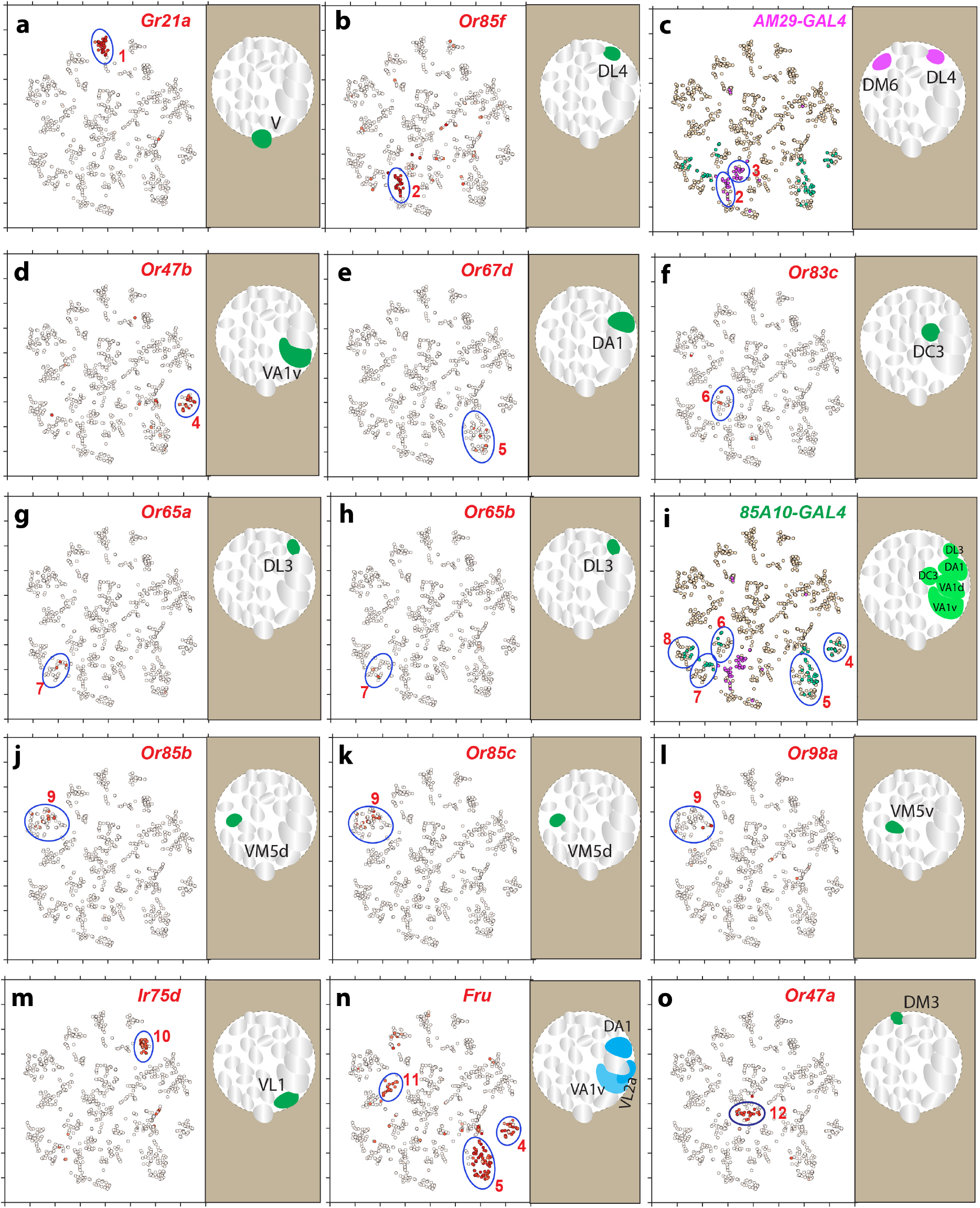
Mapping transcriptomic clusters to ORN classes. **a–o**, Clusters 1 to 12 were mapped to ORN classes using expression of different olfactory receptor genes, the gene *fruitless* (*fru*), and two different drivers, *AM29-GAL4* and *85A10-GAL4*. In each panel, a tSNE plot on the left shows the expression of one gene or one GAL4 driver, and a schematic on the right shows the ORN class(es) that are labeled by the gene or the GAL4 driver. See the color bar in Extended Data Fig. 3b for expression levels.

**Extended Data Figure 5.**
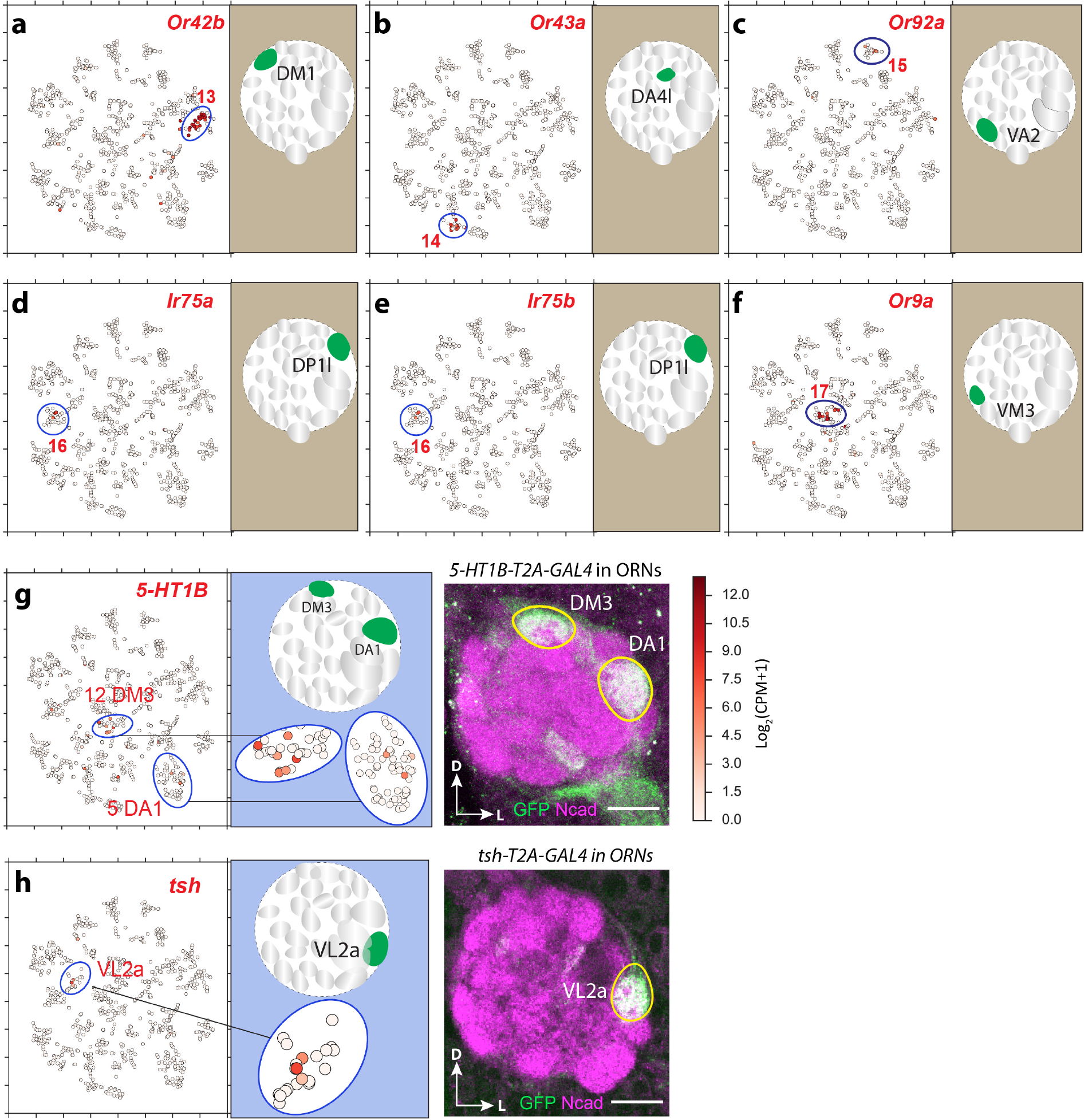
Mapping transcriptomic clusters to ORN classes and validation. **a–f**, Clusters 13 to 17 were mapped to different ORN classes using expression of different olfactory receptor genes. In each figure, a tSNE plot on the left shows the expression of an olfactory receptor gene, and a schematic on the right shows the ORN class that is labeled by the gene. See the color bar in (**g**) for expression levels. **g,** Validation of two decoded clusters that were mapped to DA1 and DM3. Confocal image shows that intersecting *ey-Flp* with *5-HT1B-T2A-GAL4* labels DA1 and DM3 ORNs at 48h APF. N-cadherin (Ncad) staining labels neuropil. Scale, 20 μm. D, dorsal; L, lateral. **h,** Validation of the decoded cluster that was mapped to VL2a. Intersecting *ey-Flp* with *tsh-T2A-GAL4* specifically labels VL2a at 48h APF. N-cadherin (Ncad) staining labels neuropil. Scale, 20 μm. D, dorsal; L, lateral.

**Extended Data Figure 6.**
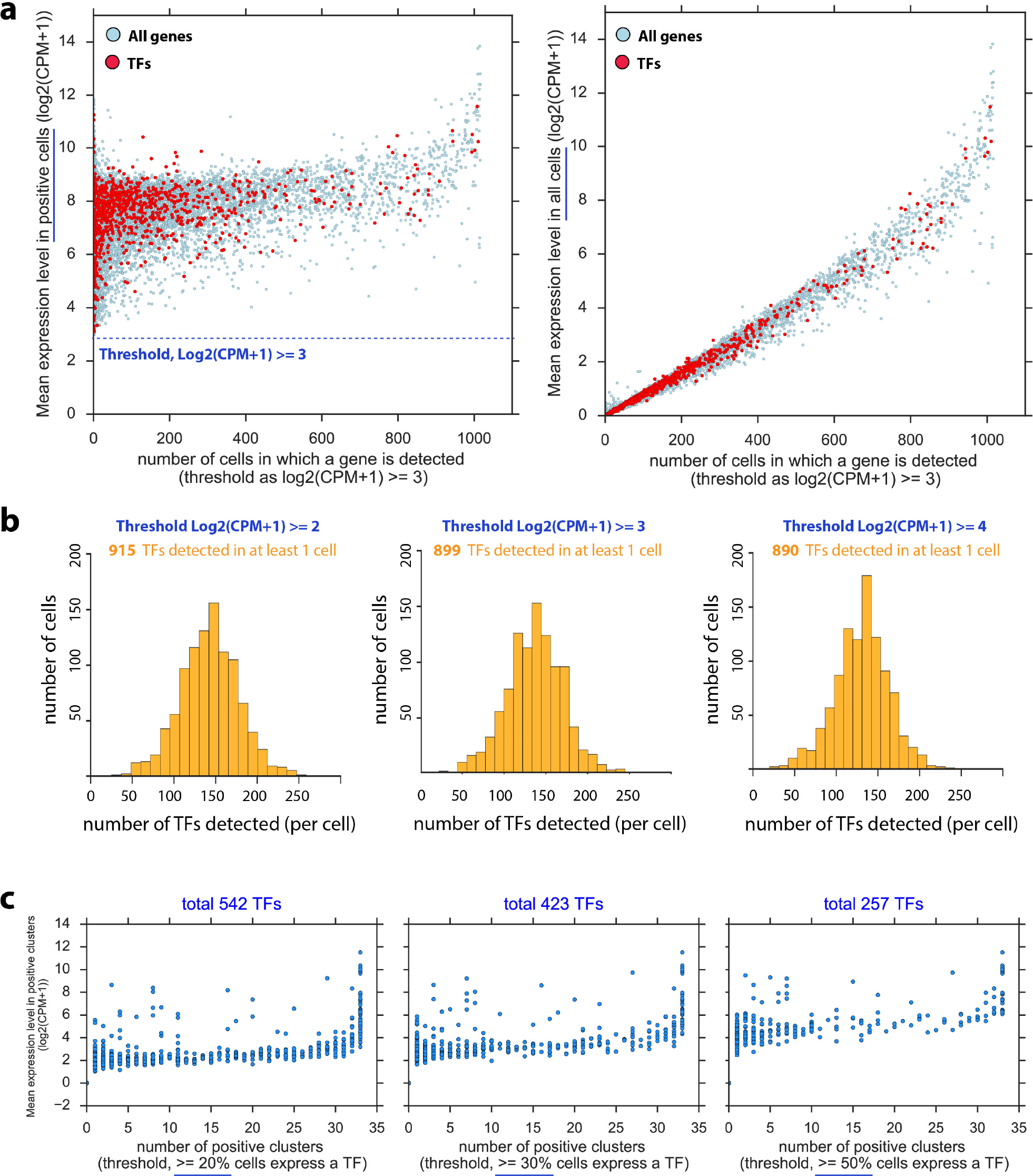
Analysis of of transcription factor (TF) expression in ORNs. **a,** Scatter plots showing number of cells in which a gene can be detected versus the mean expression level of the gene either in positive cells only (left) of in all cells (right). TFs are highlighted. A positive cell is defined as the cell expressing a gene at the level Log_2_(CPM+1) ≥3. CPM, counts per million. **b,** Similar TF distribution patterns were observed using three different thresholds of expression levels. Similar total numbers (915, 899, and 890) of detected TFs in at least one cell were obtained. ~50–250 TFs can be detected per cell, peaking at ~150. CPM, counts per million. **c,** Sparsity and expression level of the TFs among ORN transcriptomic clusters using three different percentage thresholds. Each dot is one TF. A positive cluster is defined as more than percentage threshold (20%, 30% or 50%) of cells in the cluster expressing the TF at the level Log_2_(CPM+1) ≥ 3. For each threshold, the total number of TFs that label at least one cluster is shown. While the total number of TFs expressed decreases when the percentage threshold increases, the distributions are similar. CPM, counts per million.

**Extended Data Figure 7.**
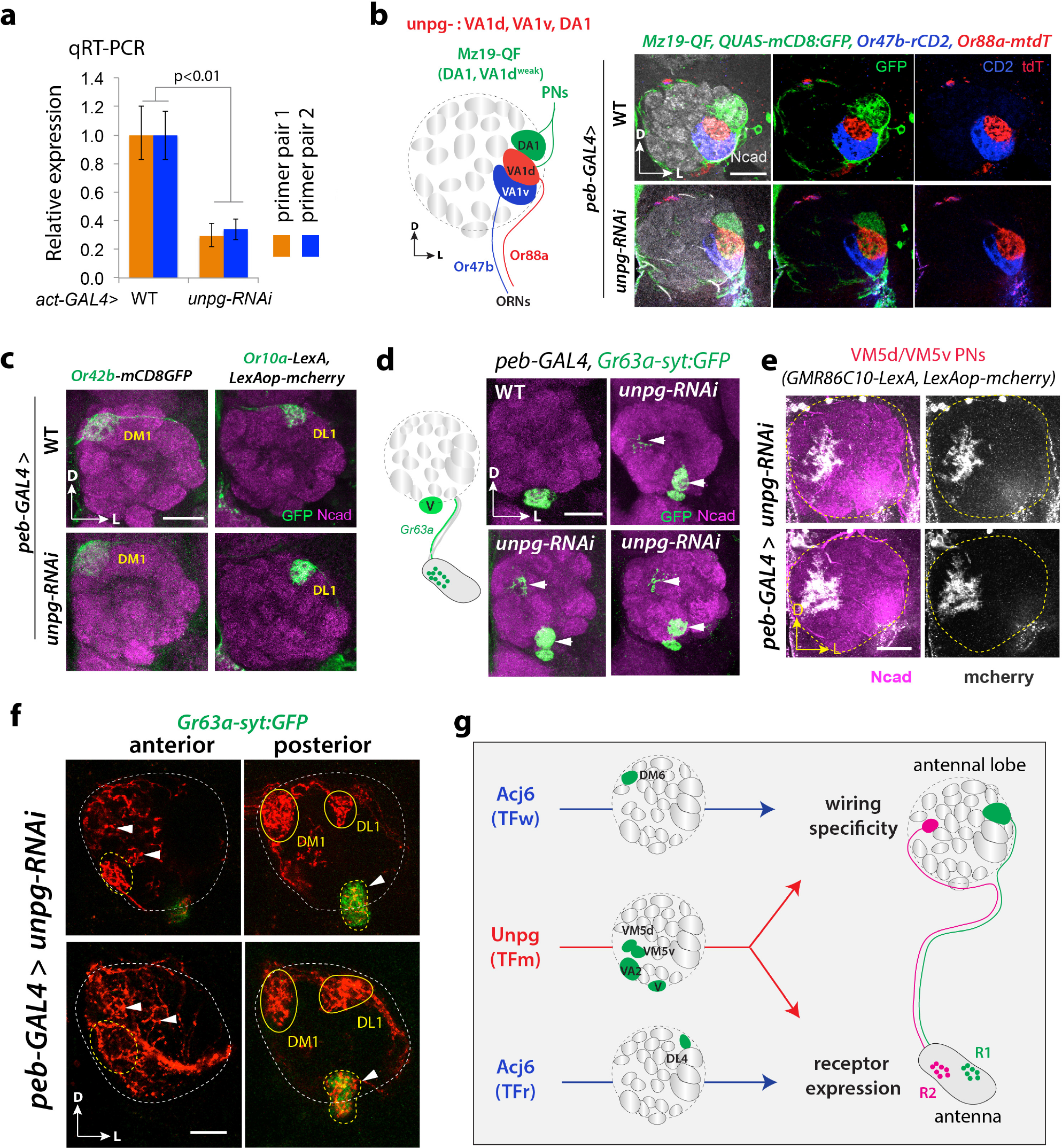
*unpg* regulates receptor expression and wiring specificity in all *unpg* + ORN classes. **a,** Quantitative RT-PCR (qRT-PCR) measurement of the knockdown efficiency of *unpg-RNAi*. *actin5C(act)-GAL4* was crossed with either *w*^*1118*^ (WT) or *unpg-RNAi*, and mRNA was extracted from pupae (*N* = 3 replicates; 5 pupae per replicate). Two pairs of primers that target to different regions of the *unpg* gene were used. Expression levels are normalized to *actin5C,* and then *WT* expressions are normalized to 1. Error bars represent SEM. *p*-value from *t* test. **b,** Confocal images showing receptor expression in control (WT) and *unpg-RNAi* flies. *peb-GAL4;UAS-dcr2* was crossed with either *w*^*1118*^ (WT) or *unpg-RNAi*. VA1d and VA1v ORNs are labeled by *Or88a-mtdT and Or47b-rCD2*. DA1 and VA1d PNs are visualized by *Mz19-QF* driven *QUAS-mCD8:GFP*. When *unpg* is knocked down, VA1d and VA1v ORN receptor expression and axon targeting, as well as DA1 and VA1d PN dendrite targeting, are all normal compared with WT control. **c,** *Or42b and Or10a* (*unpg*-negative), as well as *Gr21a and Or92a* (*unpg*-positive) ORNs are from the same ab1 sensilla in the antenna. When *unpg* is knocked down, *Or42b*+ (DM1) and *Or10a*+ (DL1) ORNs show normal receptor expression and axon targeting. This data suggests that loss of *Gr21a/Or92a* in *unpg-RNAi* flies (Fig. 4d) is not due to gross developmental defect of the ab1 sensilla. **d,** *Gr63a* and *Gr21a* are co-expressed in V ORNs. For confocal images, *peb-GAL4;UAS-dcr2* was crossed with either *w*^*1118*^ (WT) or *unpg-RNAi*, and *Gr63a-syt:GFP* is used to label V ORNs. In *unpg-RNAi* flies, V ORNs show stereotyped mistargeting (arrowheads). **e,** Additional confocal images for Fig. 4f. **f,** Additional confocal images for Fig. 4h. **g,** Summary of transcriptional regulation of receptor expression and wiring specificity in fly ORNs by two TFs, Acj6 and Unpg. Each ORN class expresses unique olfactory receptors, and sends their axons to a specific glomerulus in the antennal lobe. Acj6 acts as a TF-wiring (TFw) to regulate wiring specificity in one ORN class and a TF-receptor (TFr) to regulate receptor expression in a second ORN class. Unpg acts as a TF-master (TFm) to regulate both events. All confocal images are from adult flies, and are z-stacks covering the targeted glomeruli. N-cadherin (Ncad) staining labels neuropil (b-e). Scale, 20 μm. D, dorsal; L, lateral.

